# Single-cell Atlas Unveils Cellular Heterogeneity and Novel Markers in Human Neonatal and Adult Intervertebral Discs

**DOI:** 10.1101/2021.10.20.465219

**Authors:** Wensen Jiang, Juliane D. Glaeser, Khosrowdad Salehi, Giselle Kaneda, Pranav Mathkar, Anton Wagner, Ritchie Ho, Dmitriy Sheyn

## Abstract

The origin, composition, distribution, and function of cells in the human intervertebral disc (IVD) has not been fully understood. Here, cell atlases of both human neonatal and adult IVDs have been generated and further assessed by gene ontology pathway enrichment, pseudo-time trajectory, histology, and immunofluorescence. Comparison of cell atlases revealed the presence of two sub-populations of notochordal cells (NCs) and their associated markers in both the neonatal and adult IVDs. Developmental trajectories predicted 7 different cell states that describe the developmental process from neonatal to adult cells in IVD and analyzed the NC’s role in the IVD development. A high heterogeneity and gradual transition of annulus fibrosus cells (AFCs) in the neonatal IVD was detected and their potential relevance in IVD development assessed. Collectively, comparing single-cell atlases between neonatal and adult IVDs delineates the landscape of IVD cell biology and may help discovering novel therapeutic targets for IVD degeneration.

## INTRODUCTION

At least 30% of total adult population is suffering from lower back pain, which originates in intervertebral disc (IVD) degeneration (Andersson, 1999; de Schepper et al., 2010; Frank et al., 1996; Macfarlane et al., 1999) To date, most treatments of IVD degeneration are limited to invasive surgical interventions, such as disc replacement and spinal fusion, or pain management that is not addressing the underlying cause of IVD degeneration (Knezevic et al., 2017).

The IVD is a shock-absorbing structure that connects two adjacent vertebras and enables spine movement (Buckley et al., 2018; Vergroesen et al., 2015). The human IVD cell composition is highly heterogenous; nucleus pulposus (NP) is the inner core and the annulus fibrosus (AF) is the outer region that is confined by two endplates sandwiching the disc (Sun et al., 2020). It is widely accepted the embryonic notochord develops into the NP and that notochordal cells (NCs) disappear within the first decade of life (Séguin et al., 2018). However, the NC remnants or dormant NC can be found in the adult human IVD (McCann and Séguin, 2016; Wang et al., 2008). Additional study showed that the NCs or NC-like NP cells (NPCs) in human persist throughout life (Risbud and Shapiro, 2011). The eventual fate of human NCs is not conclusive to date, but understanding its development has considerable potential to benefit cell therapies of IVD degeneration. Animals that keep the NCs in adulthood like cats and pigs do not to exhibit disc degeneration (Hunter et al., 2003; Sheyn et al., 2019). NCs hold great potential to rejuvenate a degenerated human IVD and this potential has already been demonstrated in a mini pig model (Sheyn *et al*., 2019).

Recent advances in single-cell RNA sequencing (scRNA-seq) allow for creation of cell atlases that unveil rare sub-populations, delineate cellular heterogeneity, identify new markers, and predict developmental trajectories (Ho et al., 2021; Kelly et al., 2020; Mathys et al., 2019; Setty et al., 2019; Stark et al., 2019). Unraveling the single-cell atlas and transcriptomes of the human IVD at different ages will largely extend our understanding of its cell biology and development. To date, its heterogeneity, particularly in the early development, has not been sufficiently shown due to the limitation of traditional bulk-sequencing (Fernandes et al., 2020). Previously, multiple bulk proteomics studies delineated the general cell heterogeneity of human IVD (Rodrigues-Pinto et al., 2018; Tam et al., 2020b) and made comparison at different developmental stages (Tam *et al*., 2020b). Recent studies of single-cell atlases in non-human vertebrates resolved the cellular heterogeneity in bovine caudal IVD (Panebianco et al., 2021) and found stem cells in rat IVD (Wang et al., 2021). Another study provided single-cell analysis of human adult IVD that identified the NPC and AFC populations (Fernandes *et al*., 2020). A recent study identified chondrocytes as dominating cell population in human IVD and found small NC sub-populations (Gan et al., 2021). Yet no study has investigated cell atlases from neonatal human IVD and thus elaborated the role of NCs in development and hemostasis at a single-cell resolution.

Here, we analyze a human IVD single-cell atlas with direct head-to-head comparison between neonatal and adult human subjects to reveal the identities of all IVD populations and sub- populations of both developmental stages, discover rare cell populations and novel markers thereof, and predict the developmental trajectories from neonatal to adult IVD. We isolated cells from neonatal and adult IVDs postmortem, and analyzed them with scRNA-seq. An in-depth gene ontology term enrichment and pathway analysis was performed and developmental trajectories from neonatal to adult IVD were predicted. We further validated our findings with histology and immunofluorescence.

## RESULTS

### Human Neonatal and Adult IVD Cells are Comprised of 14 Distinct Sub-populations

Histological analysis of neonatal IVD tissue demonstrated two distinct tissue regions. Region A contained aggregated cells and loose extracellular matrix (ECM) structures and region B contained sparsely distributed cells and dense ECM tissue (Fig. 1a). Vessels and red blood cells were clearly identified in region A (Fig. 1a). In contrast, the adult IVD did not exhibit different regions. The uniform and homogenous tissue contained sparsely distributed cells whose cytoplasm was greatly larger and cell density lower than in neonatal IVD (Fig. 1b).

**Figure 1.**
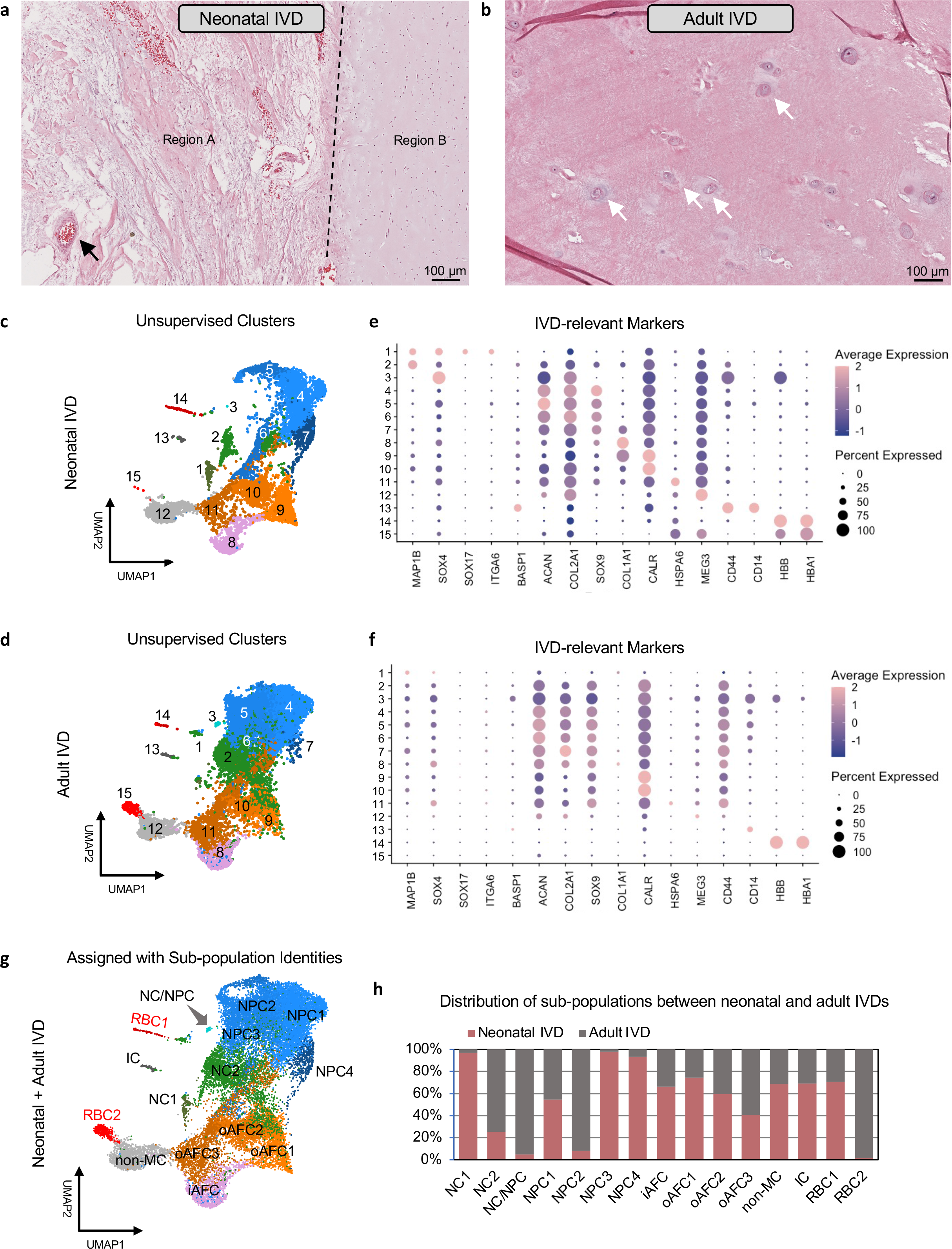
Sorting of Cells from Human Neonatal and Adult IVD of Distinct Tissue Morphologies into 15 Unsupervised Clusters Based on Their Expression of IVD Markers. H&E staining of a) neonatal IVD and b) adult IVD. The black dashed line shows the boundary separating two distinct regions in a). Black arrow points to a blood vessel. The white arrows point to NPCs. c-d) UMAP shows 15 unsupervised cell clusters that were identified in both neonatal (c) and adult (d) samples. e-f) Dot plots showing the expression level of classical IVD markers in each cluster of neonatal (e) and adult (f) IVD. Dot size indicates percent of cells expressing the genes and dot color intensity the average expression level (normalized to -2 to 2). g) Combined UMAP for all cells from neonatal and adult IVDs with clusters assigned to sub-populations based on their marker expressions in (c-d). h) Cell distribution between two ages (neonatal and adult) for each sub-population. Numbers were normalized to total cell counts for each age.

Dimensional reduction based on the unsupervised UMAP method sorted all cells into 15 clusters that could be classified as 15 cell sub-populations residing in both neonatal (Fig. 1c) and adult IVDs (Fig 1d). The expression levels of classical IVD markers in each cluster were shown as dot plot for neonatal IVD (Fig. 1c) and adult IVD (Fig. 1d). The selection of classical IVD-relevant markers includes NC markers *MAP1B* (Rodrigues-Pinto *et al*., 2018)*, SOX4* (Bhattaram et al., 2010)*, SOX17* (D’Amour et al., 2005; Sheyn *et al*., 2019)*, ITGA6 and BASP1* (Sheyn *et al*., 2019), NPC markers *ACAN* (Fernandes *et al*., 2020; Risbud et al., 2015; Tam et al., 2020a), *COL2A1* (Fernandes *et al*., 2020; Risbud *et al*., 2015; Tam *et al*., 2020a) and *SOX9* (Sheyn *et al*., 2019), AFC markers *COL1A1* (Tam *et al*., 2020b; van den Akker et al., 2017), *CALR* (Tam *et al*., 2020b) and *HSPA6* (Takao and Iwaki, 2002), as well as other previously shown markers like *MEG3 for non-mesenchymal cells* (Chen et al., 2017)*, CD44* (Schumann et al., 2015) *and CD14* (Ziegler-Heitbrock and Ulevitch, 1993) *for monocytes,* and *HBB* (Talamo et al., 2003) and *HBA1* (Pandey and Rizvi, 2011) *for red blood cells* (RBC, Fig. 1e-f). Based on these markers we annotated each cluster (Fig. 1g). The distribution of cells from neonatal vs. adult IVD in each annotated sub- population, normalized by total cells of their respective age, shows that neonatal IVD disproportionally contributed to NC1, NPC3-4, (inner) iAFC and (outer) oAFC1 clusters (Fig. 1h). Adult IVD disproportionally contributed to the NC2, NC/NPC, NPC2, and RBC2 clusters. The contribution to rest of the sub-populations was more or less evenly distributed.

### Human Neonatal and Adult Intervertebral Disc Contain Two Distinct Notochordal Cell Sub- Populations

Cluster 1 and 2 in the neonatal IVD were identified as NC sub-populations (Fig. 1c-h) based on their expression of NC markers, *MAP1B* (Rodrigues-Pinto *et al*., 2018) and *SOX4* (Bhattaram *et al*., 2010), and have been assigned to NC1 and NC2 (Fig. 2a). NC1 and NC2 were found to be distinct from each other: NC1 expressed *SOX17* and *ITGA6*, while NC2 expressed *CD44* (Fig. 2a). The NC sub-populations were also found in adult IVD (Fig. 2a). NC2 showed overwhelmingly higher cell counts than NC1 in adult (Fig. 1h and 2a). Immunostaining of the NC markers, *MAP1B* and *SOX17,* confirmed their expression in both neonatal and adult IVDs (Fig. 2b) on the protein level. These proteins showed a higher overall expression in neonatal than in adult IVD. We also observed the NC2 expressed immune-relevant marker *CD44* (Fig. 2a) and confirmed the presence of CD44 proteins in both ages (Fig. 2b).

**Figure 2.**
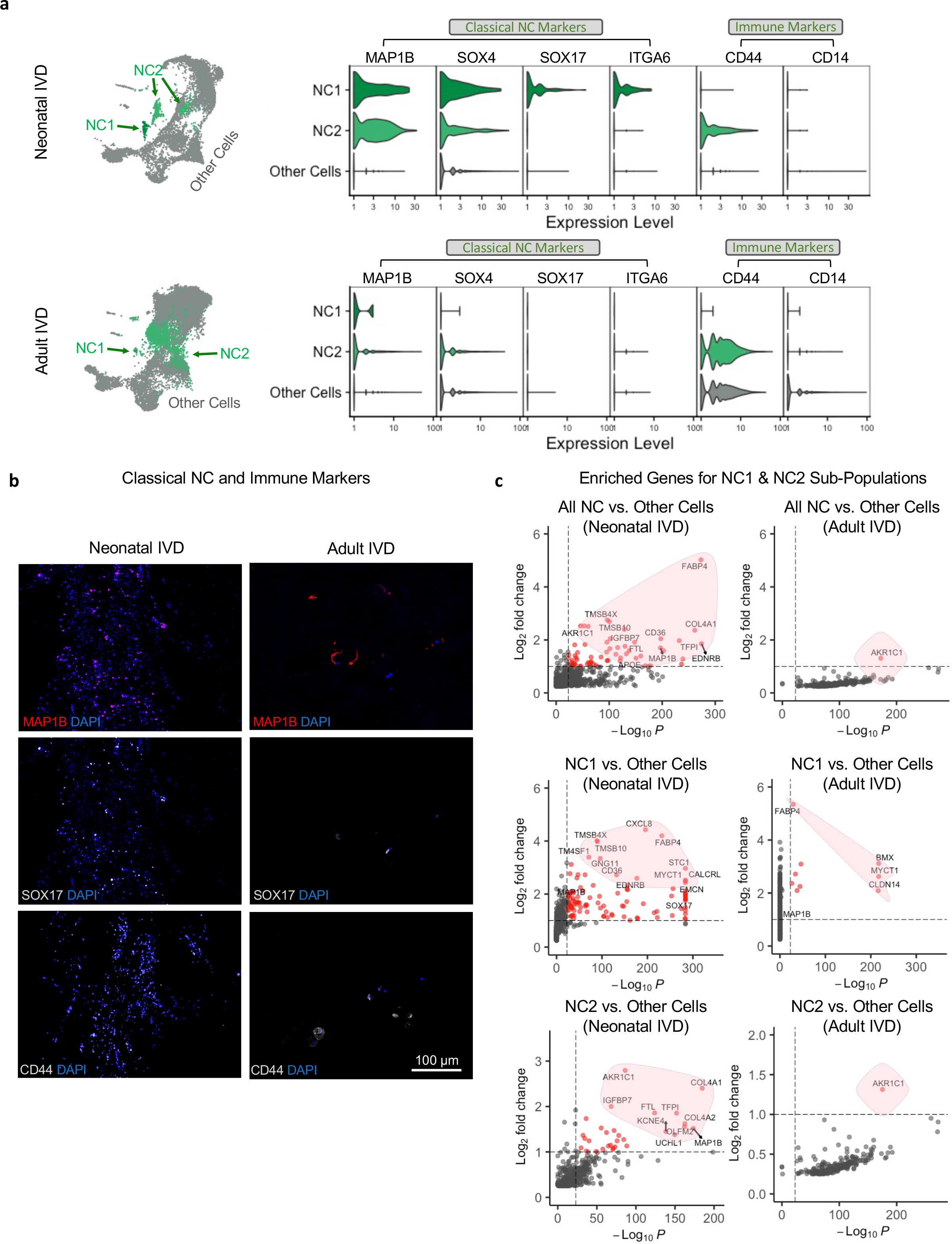
Identification of NC Sub-Populations and Their Pathway Network Analysis. **a)** The two NC sub-populations in UMAP and their expression of classical markers in both neonatal and adult IVD; two *MAP1B*+/SOX4+ NC sub-populations in the UMAP are colored green, and all other cells are colored grey; violin plots show NC expression levels of classical markers of NCs and immune cells for human neonatal and adult IVD. Other cells refer to the expression level for all cells in all clusters other than NC1 and NC2. **b)** The immunostaining of classical NC and immune markers in neonatal and adult IVD are shown. Scale bar = 100 µm; original magnification = 20x. **c)** Volcano plots showing the enriched genes for the entire populations of NC, sub-populations of NC1 or NC2, in neonatal or adult IVD. The y-axis in the volcano plots shows the -log10*p* where the *p* value was about an enriched gene expression level in a specific sub-population in a specific age (neonatal or adult) against all other cells the same age. The x-axis shows the log2FC (FC - fold change) of the expression level. The cut-off threshold was set to FC>2 and *p*<1⨉10^-24^. The enriched genes meeting the cut-off threshold are colored with red. The genes not meeting the cut-off threshold are colored with grey.

Pathway networks were further created based on the list of all enriched genes for NC1 and NC2 (Fig. S1a, b), that is, 860 genes for NC1 and 583 genes for NC2. Our results show NC1 preferentially demonstrated the activities of cancer-related pathway STAT1 and the deactivation of Brachyury (*TBXT*), Microphthalmia-associated Transcription Factor (*MITF)*, and SWI/SNF Related, Matrix Associated, Actin Dependent Regulator of Chromatin (*SMARCD3)*, which is further associated with the deactivation of connective tissue cell pathways and the collagen- relevant Glycoprotein VI (*GP6*) signaling pathway, and the deactivation of connective tissue activities (Fig. S2b). *MITF* was previously shown to be associated with osteoclast activities (Lu et al., 2010) and GP6 was shown to induce collagen deposition (Stegner et al., 2014). NC2 preferentially demonstrated stem cell pluripotency factor FGF2 (Fig. S2a) which led to the organization of cytoplasm and organization of cytoskeleton.

### *AKR1C1, APOE,* and *FABP4* are Novel Markers for Notochordal Cells in Human Intervertebral Disc

We selected several NC markers from top enriched genes that are shown in Fig. 2c. Specifically, *FABP4* is a strong marker for NC1 in both neonatal and adult IVD, *AKR1C1* is the NC2 marker for neonatal IVD, and *APOE* is a relatively weak marker but can be identified in the entire NC population (Fig. 2c). Interestingly, AKR1C1 is the only marker common to the entire NC population in adult IVD (Fig. 2c). We compared the expression levels of *FABP4*, *AKR1C1*, and *APOE* in neonatal and adult IVDs due to their possible age-dependent specificity (Fig. 3a). *FABP4* is only specific to neonatal NC1. *AKR1C1* lost certain specificity in adult IVD (Fig. 3a) but is still specific to the entire NC population and NC2 (Fig. 2c). The immunostaining of novel NC markers, *AKR1C*, *FABP4,* and *APOE,* confirmed their expression in both neonatal and adult IVDs. All markers show overlap with cell nucleus besides *AKR1C* that appears in the cytoplasm.

**Figure 3.**
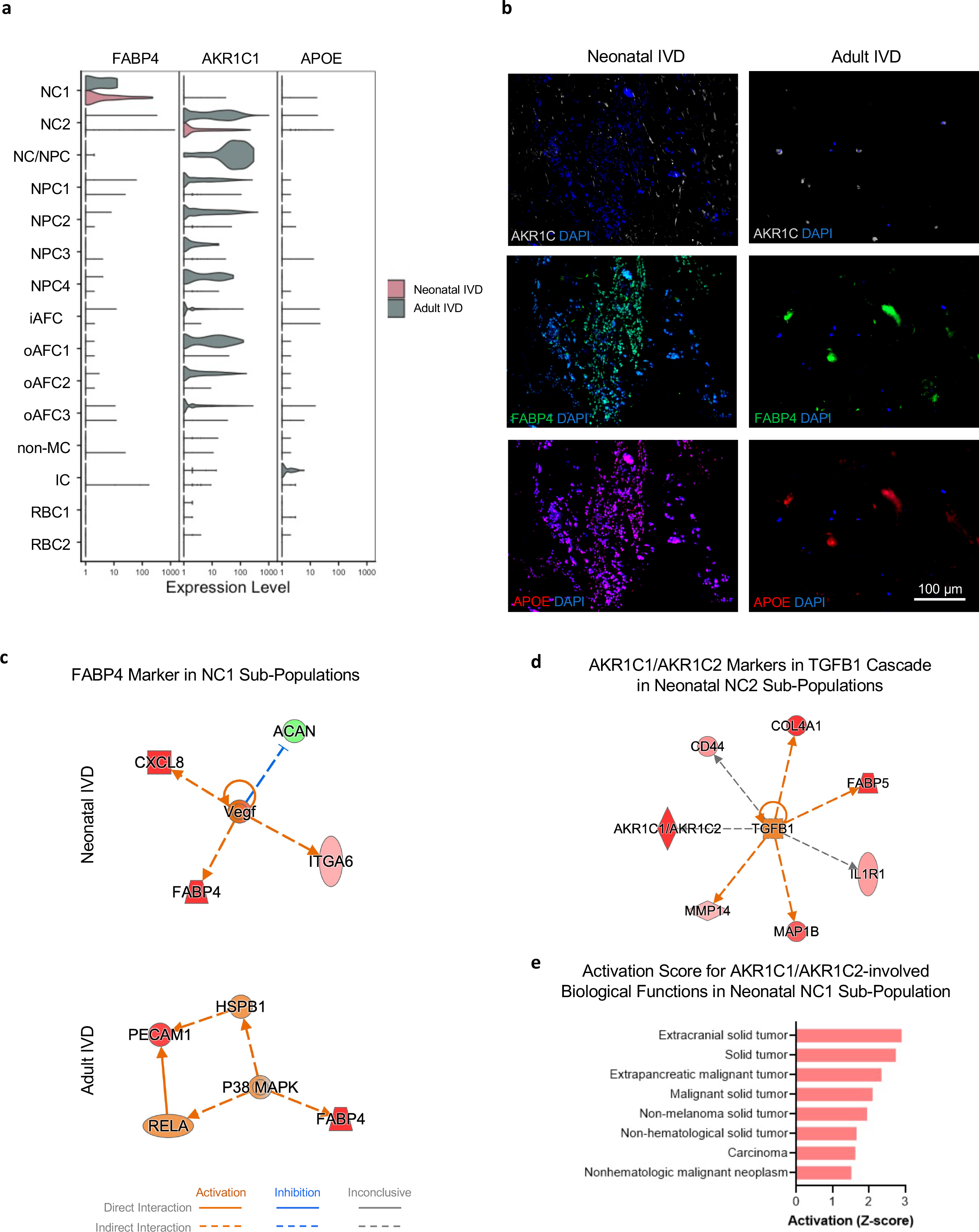
Discovery of Novel NC Markers, *FABP4, AKR1C1, APOE*, and Their Pathway/Network Analysis. **a)** Comparison of the expression level of novel top markers in human neonatal versus adult IVD. **b)** Immunostaining of novel NC markers in neonatal and adult IVD are shown. The scale bar = 100 µm and original magnification = 20x. Qiagen IPA software was used to plot the **c)** VEGF-regulated network involving *FABP4* marker in NC1 sub-population in neonatal IVD and P38-MAPK-regulated network involving *FABP4* marker in NC1 sub- population in adult IVD. **d)** TGFB1-regulated network involving *AKR1C1* marker in NC2 sub- population in neonatal IVD. **e)** Top biological functions activated by *AKR1C1*/*AKR1C2* and ranked by the activation score in NC1 sub-population in neonatal IVD.

We further investigated the role of the new markers in the regulatory networks and biofunctions. We found that *FABP4* plays a critical role in the *VEGF*-regulated network in neonatal IVD, in which the upregulation of *VEGF* leads to the activation of NC markers *FABP4* and *ITGA6*, inflammatory factor *CXCL8*, and downregulate *ACAN* (Fig. 3c). In adult IVD, however, FABP4 is activated by P38 MAPK regulator (Fig. 3c). *AKR1C1* and *AKR1C2* play a critical role in the *TGFB1* cascade in the neonatal NC2 sub-population (Fig. 3d) This network seems to be highly relevant to NCs as it also leads to the activation of classical NC markers *MAP1B* and immune-relevant *CD44* (Fig. 3d). The activation of *AKR1C1* is associated with multiple biological functions, as shown in Fig. 3e. Specifically, the upregulation of *AKR1C1/AKR1C2* activates multiple tumor and cancer-related functions (Fig. 3e). Lastly, we noticed the novel *APOE* marker was in the IL1 cascade, which also regulates *AKR1C1/AKR1C2*. (Fig. S1c).

### Nucleus Pulposus Cell Populations Contain Four Similar Sub-populations with Different Biological Functions in Neonatal Versus Adult Intervertebral Disc

NPCs expressing their classical markers, *ACAN* (Fernandes *et al*., 2020; Risbud *et al*., 2015; Tam *et al*., 2020a)*, COL2A1* (Fernandes *et al*., 2020; Risbud *et al*., 2015; Tam *et al*., 2020a), and *SOX9* (Sheyn *et al*., 2019) were detected in both neonatal IVD (Fig. 4a) and adult IVD (Fig. 4b). The entire NPC population contained four different sub-populations identified by their specific markers: NPC1 identified by *OGN*, NPC2 identified by *MMP13*, NPC3 identified by *COL9A1*, and NPC4 identified by *DPT*. The clusters 4-7 (Fig. 1c-d) thus have been designated as NPC1-4 respectively. Cluster 3 (Fig. 1c-d) partly expresses NC markers (SOX4+ but no MAP1B) and partly expresses NPC markers (COL2A1+ but no ACAN and SOX9), and thus it has been designated as NC/NPC. It may represent a transition state of cells from NC to NPC or a reservoir of NP progenitors. The expression of the NPC markers is shown in violin plots (Fig. 4a-b). However, four NPC sub-populations from the same age are barely distinguishable (Fig. 4a, b). The sub- population markers are not very specific as the gene expression distributed on the UMAP did not always match the clustering on the UMAP (Fig. 4a, b). The four NPC sub-populations may have similar transcriptomic profiles. The AFC marker levels were used as controls (Fig. 4a, b) and their expression was found very low among all NPCs in both the neonatal and adult IVD, which confirmed the NPC clusters are not overlapping with AFCs. Only NPC4 in neonatal IVD expressed AFC marker *COL1A1* (Fig. 4a).

**Figure 4.**
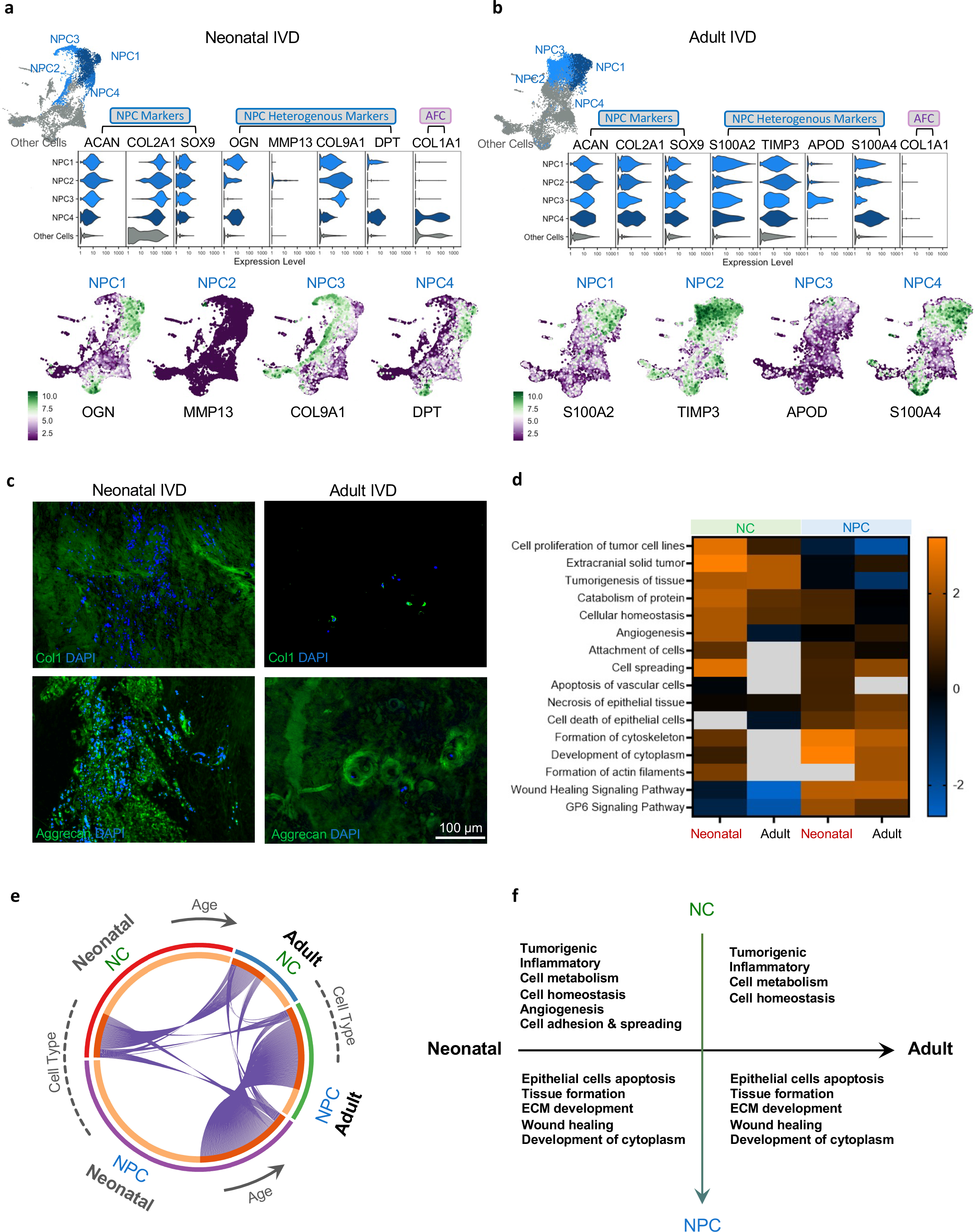
Identification of 4 Similar NPC Sub-Populations and Comparison of Their NPC Gene Expression, Pathways, and Biofunctions with NC Populations in Neonatal and Adult IVD. **a-b)** Violin plots showing differences in the expression levels of NPC markers in four NPC sub-populations (NPC1-4) identified among the *ACAN+/COL2A1+/SOX9+* NPC populations in UMAP (colored in blue) in both neonatal (a) and adult IVD (b). AFC markers were shown for comparison. Expression levels of each NPC subtype marker were projected via UMAP. **c)** Immunostaining of extracellular matrix (ECM) proteins in neonatal and adult IVD is shown. Scale bar = 100 µm; Original magnification = 20x. **d)** Heatmap shows biological functions enriched from all DEGs of NCs or NPCs in neonatal or adult IVD. Orange indicates activation, deactivation, and grey data that were not detected or did not pass the filtering. **e)** Circos plot showing the overlap between four different enriched gene lists: neonatal NCs, adult NCs, neonatal NPCs, and adult NPCs. Each purple line links the same gene expressed in both enriched gene list. Red and orange show the shared and unique genes. **f)** Descriptive summary terms on the biofunctions enriched from each list.

NPCs are critical to the hemostasis and deposition of ECM (Frapin et al., 2019). We observed that the age greatly affected the critical ECM proteins (Fig. 4c). Figure 4c shows the abundant presence of collagen type 1 (*COL1A1*) in the extracellular space and aggrecan in the cytoplasm in the neonatal IVD. The signal of intracellular *COL1A1* appears to be weak in neonatal IVD (Fig. 4c). Figure 4c shows the opposite trend for adult IVD: the staining of *COL1A1* in the extracellular space of neonatal IVD is negligible, but the staining in the cytoplasm is strong; the staining of aggrecan in the extracellular space is strong, but weak in the cytoplasm (Fig. 4c).

### Notochordal Cell and Nucleus Pulposus Cell Populations Exhibit Different Gene Ontology Enrichments, Pathways, and Regulators

NC and NPC populations were compared in both neonatal and adult IVDs (Fig. 4d-f). In this analysis, NC1-2 were combined as NCs, and NPC1-4 were combined as NPCs (Fig. 4d). NC populations were preferentially enriched to cancer and tumor activities, cell metabolism, cellular homeostasis, angiogenesis, and cell adhesion and spreading, whereas NPC populations were preferentially enriched to cytoskeletal tissue formation, the development of cytoplasm, wounding healing and ECM development (Fig. 4d). We also showed top-scored key regulatory networks in neonatal NCs and adult NPCs in Fig. S1d. Several inflammatory and tumorigenic factors and pathways (*IL1B, VEGF*, *HGF*) were upregulated in NC (Fig. S1d). In NPCs, upregulation of the osteogenic factor *BMP2* (Cai et al., 2021), osteoclast regulator *MITF* (Lu *et al*., 2010), DNA-binding transcription factor-related *BCL6*, NPC and skeletal development marker *SOX9* (Bi et al., 1999; Sheyn *et al*., 2019; Zhou et al., 2006) were detected (Fig. S1d). This trend seems to be age-independent except for *BCL6*, which was not detected in the adult IVD (Fig. S1d). Different cell types (NC vs. NPC) demonstrated hugely distinct transcriptomes, but the differences between the ages (neonatal vs. adult) was much smaller, as summarized in the Circos plot (Fig. 4e), where the overlap of expressed genes mostly occurred among the same cell types. The summary of enriched functional terms is presented in Fig. 4f.

### Annulus Fibrosus Cells in Neonatal IVD are Highly Heterogenous, and their Expression of Collagen-Relevant Genes is Location-Dependent

*COL1A1* is the most common ECM protein secreted by AFCs (Tam *et al*., 2020b; van den Akker *et al*., 2017). *CALR* is a marker for the outer AF (Tam *et al*., 2020b) and heat shock proteins (*HSPs*, e.g. *HSPA6*, *HSPA1A*, etc.) are markers for the most outer layer of the AF (Takao and Iwaki, 2002). Thus, the *COL1A1, CALR,* and *HSPA6* were used to identify AFC populations (Fig. 5a, b). Cluster 8 (Fig. 1c-d) has been classified as the inner AFC sub-population (iAFC), due to its expression of *COL1A1* (Fig. 5a, b). The clusters 9-11 (Fig. 1c-d) have been assigned to the outer AFC sub-population (oAFC1-3) respectively, due to their expression of *CALR* (cluster 9, 10) and *HSPA6* (cluster 11, Fig. 5a, b). We detected a high degree of heterogeneity among neonatal AFC populations (Fig. 5c). We identified *LGALS1, HES1, HERPUD1,* and *DNAJB1* as AFC sub- population markers for iAFC and oAFC1-3. Moreover, several types of collagens were also detected in neonatal IVD, specifically *COL1A1, COL3A1, COL5A1, COL5A2, COL6A3, COL12A1* (Fig. 5d). Strikingly, their expression showed a decreasing trend assuming the location of subtypes follows the order of iAFC, oAFC1, oAFC2 and oAFC3 from inner core to the outer region (Fig. 5d). A similar trend was observed in the enriched biofunctions and key pathway analyses (Fig. 5e) across the neonatal iAFC and oAFC populations. As part of the pathways listed in Fig. 5e, we show the mechanistic scheme of Unfolded Protein Response (UPR) signaling pathway and the relevant gene expression in oAFC2 in Fig. S2. The adult IVD did not exhibit similar trend of collagen-relevant gene expression (Fig. S3). The illustration in Fig. 5f shows the postulated structure transition from neonatal to adult IVD based on these spatially dependent gene expression patterns.

**Figure 5.**
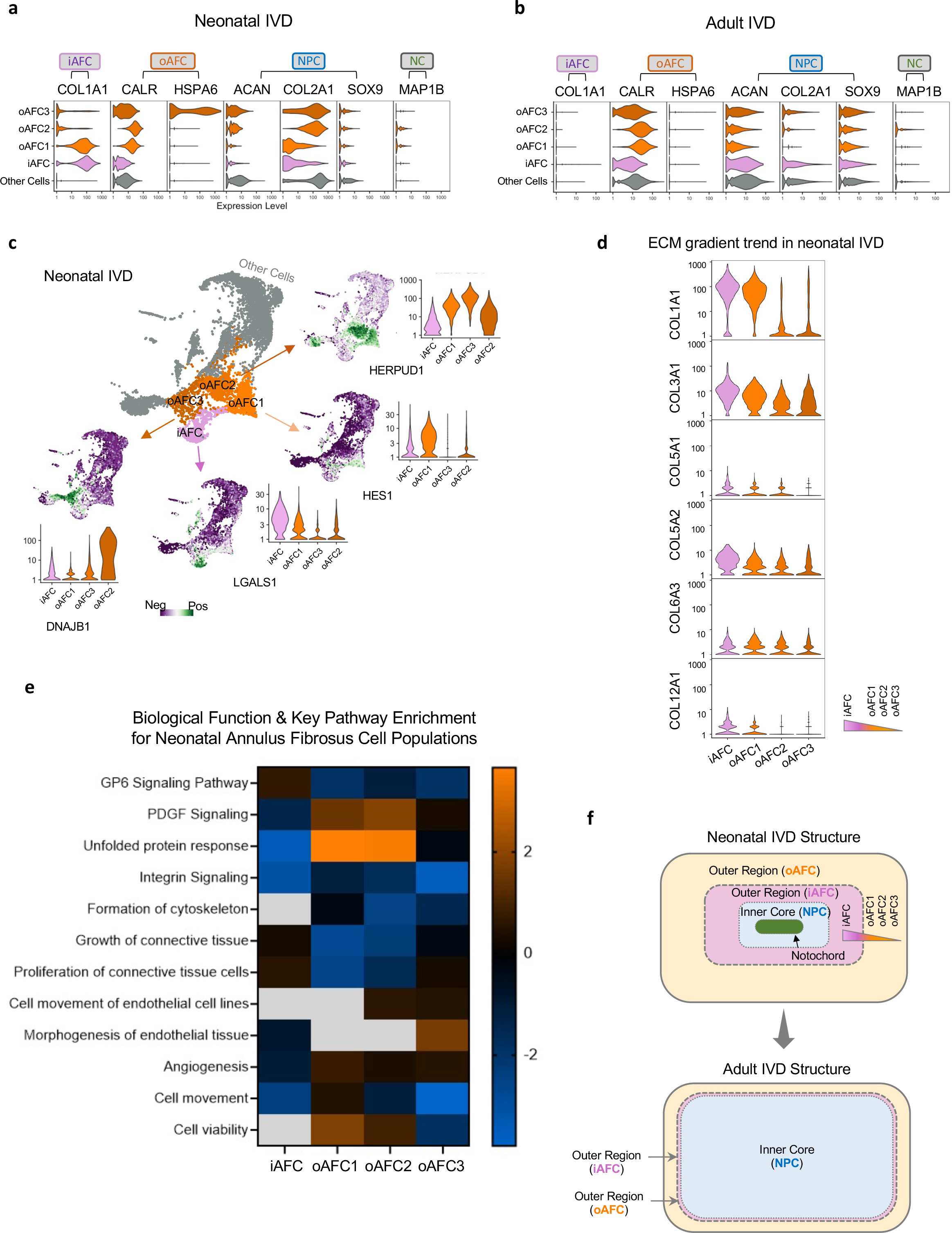
Heterogeneity of Annulus Fibrosus Cells in Human Neonatal and Adult IVDs. **a-b)** Expression levels of classical AFC markers (*COL1A1, CALR, HSPA6*), classical NPC markers *(ACAN, COL2A1, SOX9*) and classical NC markers (*MAP1B*) are shown for comparison, for both (**a)** neonatal IVD and **(b)** adult IVD. **c)** The AFC populations showed strong heterogeneity in neonatal IVD by having 4 distinct subtypes and their heterogenous markers. We demonstrated the expression distribution projected on UMAP and their quantitative expression levels. **d)** Among the 5 heterogenous sub-populations detected in **(c)**, the neonatal AFCs exhibited a decreasing trend of expression levels of ECM-relevant, collagen-producing genes from inner core NPCs to the outer region AFCs following the order of iAFC, oAFC1-3. **e)** The biological function and key canonical pathway enrichment results followed the same gradient. **f)** Predicted scheme for spatial distribution of the 4 AFC populations, along with inner NPC core and notochord, in neonatal IVD, as compared with the classical IVD structure in adult human.

### Resolved Single-cell Atlas for Human Neonatal and Adult IVD with Clusters Assigned to Sub- populations

We have assigned cluster 12 into non-mesenchymal cells (non-MC) due to their *MEG3* expression, a marker for non-mesenchymal, tumor-suppressing cells (Chen *et al*., 2017). Cluster 13 is a satellite cell population on the UMAP (Fig. 1c-d). It shared the *CD44* expression as NC2 but lacked all other NC markers (Fig. 1c, d). Thus, we have assigned cluster 13 to immune-like cells (IC) due to its expression of immune cell marker CD14 (Ziegler-Heitbrock and Ulevitch, 1993) and CD44 (Schumann *et al*., 2015) (Fig. 6a). Cluster 14 and 15 have been assigned to RBC2 an RBC1 due to its expression of hemoglobin markers *HBB* (Talamo *et al*., 2003) and *HBA1* (Pandey and Rizvi, 2011) (Fig. 6a). In addition to classical markers that were used to assign cell identities, Figure 6a also includes novel markers we found for NC, AFC and NPC populations. A heatmap of the expression levels of top enriched genes for all sub-populations shows the high specificity of our enriched genes and the high cellular heterogeneity in IVD structures (Fig. S4). Collectively, the single-cell atlas of human IVD heterogeneously comprised of NC, NPC, AFC, IC, non-MC, and RBC populations, which can be more precisely classified into 15 different subtypes: NC1-2, NC/NPC, NPC1-4, iAFC, oAFC1-3, non-MC, IC and RBC1-2 (Fig. 6a).

**Figure. 6.**
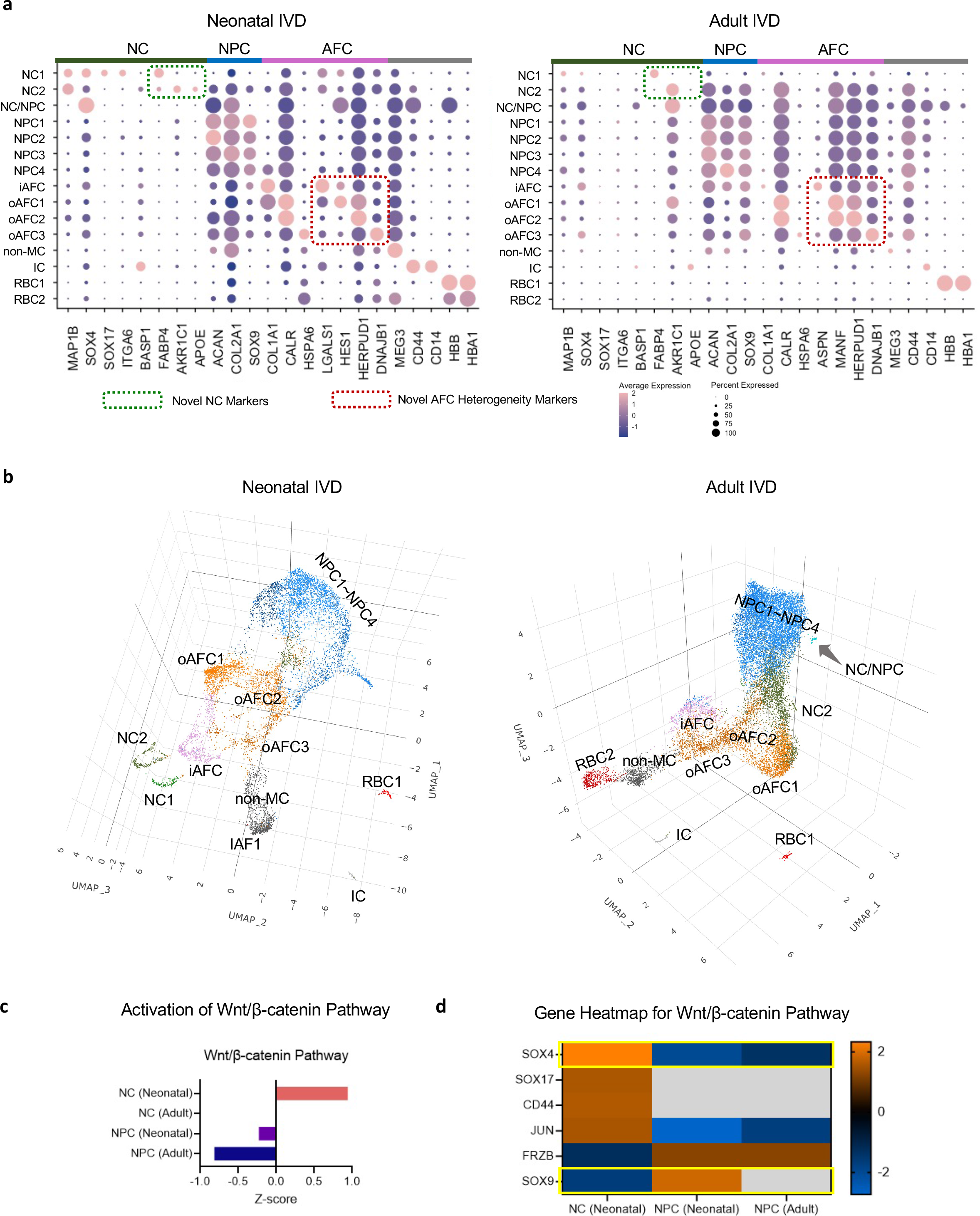
Complete Single-cell Atlases of Human Neonatal and Adult IVD. **a)** Complete dot plots of marker expression of all sub-populations in the neonatal and adult IVD, including classical markers and novel markers found in the present study. The percent expressed is shown by the size of the dot, and the average expression level (normalized to -2 to 2) is shown by the color scale. **b)** Complete single-cell atlases visualized with 3D UMAP method, for both neonatal and adult IVD. The gene markers in the green rectangle are the novel markers found in the present study. **c)** Activation Z-score rated by Qiagen IPA software showing the canonical Wnt/β-catenin pathway has been differentially regulated in NCs and NPCs in neonatal or adult IVD. **d)** Wnt/β- catenin pathway key molecules’ expression levels are shown in the heatmap. The orange indicates the activation, the blue indicates the deactivation, and the grey indicates the data either was not detected or did not pass the filtering.

The fully resolved single-cell atlases were plotted in three-dimensional (3D) UMPAs (Fig. 6b) for a more accurate and intuitive visualization. In the 3D atlas, NC1, NC2, IC, RBC, and non-MC visually appear to be away from the main continents. The NPC sub-populations (NPC1-4) tended to cluster together with vague boundaries whereas AFC sub-populations (iAFC, oAFC1-3) are distant from each other with clear boundaries.

One of the key pathways involved in the IVD is the Wnt/β-catenin pathway (Kondo et al., 2011). We found that the Wnt/β-catenin pathway to be activated in neonatal NC but downregulated in neonatal and adult NPCs (Fig. 6c). We further expanded the key genes in the Wnt/β-catenin pathway and showed the expression of *SOX4* regulator (also the NC marker) was upregulated in neonatal NCs compared to neonatal and adult NPCs, whereas another regulator gene, *SOX9,* was downregulated in neonatal NCs compared to neonatal NPCs (Fig. 6d).

### The Pseudo-time Trajectory Predicted Five Developmental States from Neonatal Cells into Adult Nucleus Pulposus Cells with Some Notochordal Cells Preserved Throughout Adulthood

The trajectory, color-coded by age, shows that neonatal cells occupy two branches and share one branch with the adult cells (Fig. 7a). Adult cells occupy the other two branches (Fig. 7a). ECM gene expression has changed over pseudo-time. *COL1A1* expression decreased over pseudo- time, while *COL2A1* decreased in the early stage, but increased in the later stage. *ACAN* expression followed a swinging increase-decrease-increase pattern (Fig. 7b). *SOX4,* a positive regulator of the Wnt/β-catenin canonical pathway (Sinner et al., 2007), decreased over pseudo- time, but *SOX9,* a negative regulator (Sinha et al., 2021), increased (Fig. 7c), which is consistent with Fig. 6d.

**Figure 7.**
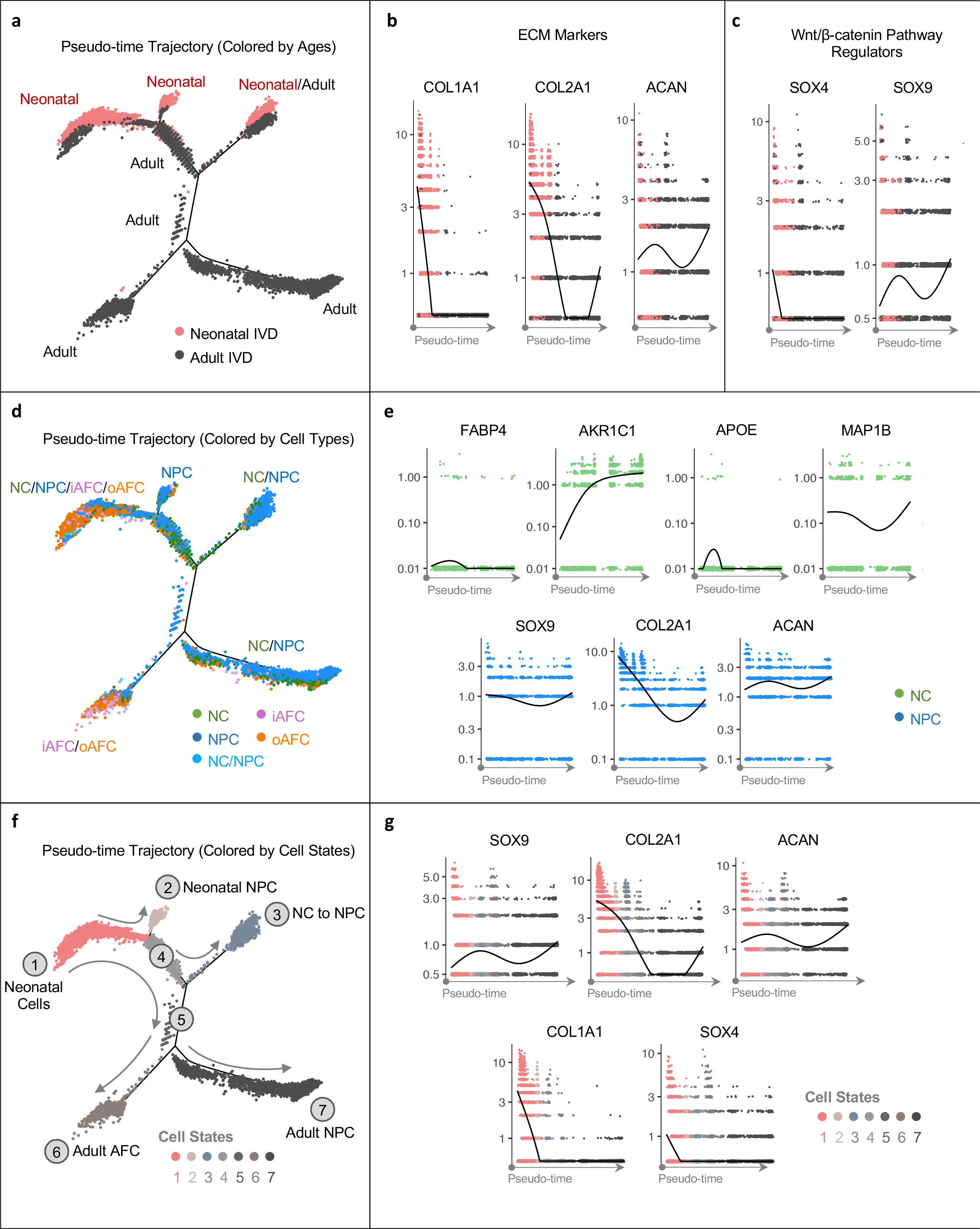
Pseudo-time Trajectories Predicted IVD Cellular and Molecular Transition During Development. **a)** Cell trajectories predicted based on 500 top genes enriched in each sub- population and color-coded based on different ages. **b-c)** The pseudo-time transition of **b)** ECM markers and **c)** Wnt/β-catenin canonical pathway regulators. **d)** Cell trajectories color-coded by different cell types, including NC, NPC, iAFC, and oAFC, and **e)** the change of classical and novel NC markers in NCs over pseudo-time, and change of classical NPC markers in NPCs over pseudo-time. **f)** Cell trajectories color-coded by different cell states. Solid grey line shows the predicted developmental direction, and the dashed grey line shows the possible but inconclusively predicted developmental direction. **g)** Pseudo-time transition of molecules of interest are displayed and colored with cell states.

NC populations are mainly positioned in the proximity to NPCs. NCs were present along the trajectory from the very early neonatal stage towards the later stage and eventually aggregate around NC/NPC in the adult branch (Fig. 7d). The two adult IVD-prevalent branches show very distinct cell compositions, one is mainly composed of NCs and NPCs, the other one is mainly composed of AFCs (Fig. 7d). The expression of novel and classical NC markers in the NC populations changed over pseudo-time, as shown in Fig. 7e. The expression of NPC markers in NPC populations also changed over pseudo-time (Fig. 7e). Specifically, *COL2A1* decreased most dramatically over pseudo-time, but bounced back in the end, *SOX9* and *ACAN* maintained relatively stable levels in NPC populations.

We found 7 cell states along the trajectory (Fig. 7f). The annotation of each state depends on the trajectories of age (Fig. 7a) and cell types (Fig. 7d). State 1 has been annotated as neonatal cells due to its mixed cell composition originated in neonatal IVD. State 2 has been annotated as neonatal NPC, as it is still in neonatal stage, but mostly consists of NPCs. State 3 has been annotated as NC to NPC, because it is comprised of both neonatal and adult cells and consists of both NCs and NPCs. State 4 and State 5 are two intermediate stages that contain NPCs, NCs, and AFCs. State 6 has been annotated as adult AFC, because it is dominated by adult cells, mainly of iAFCs and oAFCs. State 7 has been annotated as adult NPC, because it is dominated by adult NPCs with a small quantity of NCs and AFCs. The direction of trajectory is determined by the direction of pseudo-time. The change of expression levels of key molecules during the developmental process are also shown over pseudo-time in Fig. 7g, including *SOX4*, *COL2A1*, *ACAN*, *COL1A1* and *SOX4*. Collectively, state 1 might indicate an early developmental stage whereas state 2 and 3 may indicate the process of NCs developing into NPCs which may occur during the neonatal stage. State 6 might indicate that AFCs are present in both ages during the development, but they may develop from neonatal AFCs into adult AFCs. State 7 may indicate the development of large quantity of NPCs in adult that are likely originating from neonatal NCs. It also indicates the continuing presence of NCs in adult IVDs.

## DISCUSSION

This study provides the first direct comparison of cell atlases at single-cell level between neonatal and adult human IVDs. With the help of scRNA-seq technology, we discovered *FABP4*, *AKR1C1*, and *APOE* as novel markers for human NCs. Our data also supported the previous assumption of the presence of NCs in the adult. It also models the developmental process from neonatal IVD, composed of various cell types, into adult IVD, composed mainly of NPCs and AFCs. Lastly, AFCs in neonatal IVD were found to be highly heterogenous with a strong gradient profile that most probably correlated with spatial distribution.

This study identifies *FABP4*, *AKR1C1*, and *APOE* as highly specific markers for NC populations in the human neonatal and adult IVD (Fig. 3a, b). Although our network analysis demonstrated both *AKR1C1* and *FABP4* to be associated with tumorgenicity and inflammation (Fig. 3c-e), these genes have also been associated with cell metabolism and may be therefore linked to increased cell activity and biosynthesis in early development. For example, *AKR1C1* is an enzyme that catalyzes NAPDH-dependent reductions (Zeng et al., 2017) and is responsible for hormone secretion (Marín et al., 2009). *FABP4* is expressed in embryonic cartilage and bone tissues (Urs et al., 2006). In addition, *AKR1C1* and *FABP4* also mark the inflammatory response in early-stage development, as shown by *VEGF*-regulated *FABP4* in neonatal NC1 sub-population that also activated CXCL8 (Fig. 3c). Similarly, *AKR1C1* and *APOE* are involved in another *IL1*-acitivated, inflammation network (Fig. S1c). In previous research, the IL1 cascade was already considered an inflammatory and catabolic marker for adult and degenerative human IVD (Johnson et al., 2015). To our knowledge, its occurrence in neonatal IVD has not been previously reported. Noteworthy, *FABP4* appears to be a robust NC1 marker in both neonatal and adult IVD. Although AKR1C1 can be used as NC2 marker, it is less specific in adult NC2 (Fig. 3a). This may indicate a reduced NC cell metabolism in the adult IVD compared to neonatal.

Our results show that *APOE* is highly specific to NCs in neonatal IVD (Fig. 2c), in contrast to a previous study in mouse IVD that demonstrated *APOE* to be expressed in both AFCs and NPCs (Zhang et al., 2013). Differences in findings may be due to the different species investigated.

Isolating and identifying rare NC populations in adult IVD is a challenging task due to their small quantity. Our single-cell atlas detected rare NC populations in adult IVD expressing *MAP1B* marker, unveiled two distinct NC sub-populations, NC1 and NC2 in neonatal IVD, and confirmed their presence in adult IVD (Fig. 2a). To our knowledge, only one recent single-cell study by Gan et al. reported a NC population in adult IVD. However, they used *TBXT* to identify NCs, whereas we used *MAP1B* (Gan *et al*., 2021). Other studies, including proteomics (Tam *et al*., 2020b) and scRNA-seq (Fernandes *et al*., 2020) of human adult IVD, reported the presence of NPCs and AFCs, but did not report NC populations. The differences in findings might be a result of different samples analyzed or analysis techniques (Fernandes *et al*., 2020; Tam *et al*., 2020b). Partly in line with our study, Richardson et al. demonstrated the presence of a sub-population with NC-like phenotype but NPC-like morphology in adult IVD, using qPCR and immunohistochemical analysis (Richardson et al., 2017). In contrast to our findings (Fig. 2a), they detected an expression of *CD24* and *TBXT*. In a later study, Rodrigues-Pinto et al. from the same group reported the pre-selected *CD24+* cells contained NCs expressing *MAP1B* (Rodrigues-Pinto *et al*., 2018), in fetal IVD. To our knowledge, our study is the only report demonstrating two distinct *MAP1B*+ NC sub- populations in adult IVD. Differences in results between Richardson et al., Rodrigues-Pinto et al. and our study might be due to several reasons: The work by Richardson et al. focused on the *CD24*+ sub-population as NC-like phenotype and observed their NP-like morphology (Richardson *et al*., 2017), while our study is focused *MAP1B*+ NC cells, since interestingly CD24+ cells were not identified by our analysis as a distinct cluster. Therefore, their *CD24*+ sub-populations may be a different, more heterogenous population across our clusters. Rodrigues-Pinto et al. sorted and specifically analyzed *CD24*+ cells (Rodrigues-Pinto *et al*., 2018), while our study did not include any pre-selection process and instead unbiasedly analyzed all cells. Lastly, our scRNA-seq technique identified two NC populations, which we divided into NC1 and NC2 (Fig. 2a), which were not detected in previous studies (Fernandes *et al*., 2020; Gan *et al*., 2021).

Historically, NPCs are considered to be derived from NC lineage (Risbud et al., 2010). We confirmed the close lineage relations between NCs and NPCs and further unveiled the transition process. Firstly, our developmental trajectories predicted that NCs started to develop into NPCs even in the neonatal stage, and its continuing development resulted in a large quantity of NPCs in adult IVD (Fig. 7f). To the best of our knowledge, no previous study predicted the developmental trajectories crossing ages (early vs. late development) and cell types (NC versus NPC). One recent study predicted the differentiation of NPCs alone based on pseudo-time trajectories without specifying the actual ages (Gan *et al*., 2021), however, it did not include neonatal IVD that is rich in NCs. Secondly, we found significant distinctions between neonatal NC and adult NPC populations at molecular, cellular, and tissue levels. We showed that neonatal NCs have some tumor-like properties and are involved in tissue growth (Fig. 4d-f), as suggested by their expression of relevant genes (*IL1B*, *VEGF*, *HGT*, Fig. S1d) and enriched functions (Fig. 4d). In contrast, adult NPCs appear to be more mature, with closer association to fully differentiated cells and developed cytoskeletal tissues (Fig. 4d-f), and they expressed *BMP2* and *SOX9* (Fig. S1d). Our findings are in line with a previous IVD single-cell transcriptomic analysis showing *BMP2* and *SOX9* expression in some adult sub-populations (Gan *et al*., 2021). Changes were also observed in cell morphology. The adult NPCs have large cytoplasm, as opposed to the small neonatal NPCs (Fig. 1a, b). This is in line with the enriched function of development of cytoplasm for adult NPCs (Fig. 4d). The large cytoplasm of NPCs (Fig. 1b) contradicts a previous report stating that NCs are larger than NPCs (Rodrigues-Pinto *et al*., 2018). We found the matrix also changed from a fibrous, less-developed appearance to a more mature cartilage-like, (Fig. 1a, b), as well as the ECM gene expression transformed from a collagens-rich (*COL1A1* and *COL2A1*) to aggrecan-rich (*ACAN*) in the later stage (Fig. 7b). It is known that *COL2A1* ensures the removal of notochord during the development (Aszódi et al., 1998). Thus, the decrease of *COL2A1* (Fig. 7b, g) may mark the successful removal of notochord and completion of NC to NPC transition. These distinctions at molecular, cellular, and tissue levels highlight the changes occur in neonatal NC during their development into adult NPC. Though most NCs differentiated into adult NPCs, some NCs kept their identity throughout the entire trajectory (Fig. 7d). The NC quantity is significantly decreased in adult (NCs constituted 7.5% of total cell counts in neonatal IVD and 17.5% in adult IVD, data not shown). Previous studies were not conclusive on their presence (Fernandes *et al*., 2020; McCann and Séguin, 2016; Risbud and Shapiro, 2011; Séguin *et al*., 2018; Wang *et al*., 2008). It should be noted that NC1 almost disappeared in adult IVD but NC2 percentage significantly elevated (Fig. 1h). This indicates that NCs became more involved in ECM interactions and tissue developments but maintained robust pluripotency during the development. (Fig. S1a and S1b). Our study shows that the NCs kept expressing NC markers, *MAP1B and AKR1C1*, in later pseudo-time and even elevated in the very late stage (Fig. 7e). Therefore, the NCs may have preserved their capability to differentiate into NPCs during their continuing presence of NCs throughout the adulthood.

The high heterogeneity of AFCs found in neonatal IVD is in line with recent studies that also described spatially defined AFC populations divided into inner and outer regions (Shi et al., 2015; Tam *et al*., 2020a). Our study presented a very detailed classification of AFCs into one sub- population to inner AF regions (iAFC), and three distinct sub-populations in the outer AF regions (oAFC1-3), which has not been reported in previous relevant studies (Shi *et al*., 2015; Tam *et al*., 2020a).

A striking trend of decreasing expression levels of collagen-associated genes (Fig. 5d) appeared when the order of the five AFC sub-populations from neonatal IVD was re-arranged from the inner region to the outer region to iAFC, oAFC1, oAFC2, oAFC3 (Fig. 5d). Interestingly, another proteomics study by Tam et al. (Tam *et al*., 2020b) showed ECM proteins in 16- to 17-year-olds increased from the inner NP to the outer AF region of the IVD, which seems to be contradictory to our findings of the decreasing expression of collagen-producing genes (Fig. 5d). The protein content and RNA level may not necessarily be aligned. In line with Tam et al., we found GP6 signaling pathway relevant to cell-ECM interaction show gradient of expression from inner AF to outer AF (Fig. 5e). The oAFC and iAFC have distinct gene expression profiles possibly due to their different spatially defined roles. AFCs are in a position that connects endplate and inner NP core, as suggested in Fig. 5f. The promoted Unfolded Protein Response (*UPR*) we detected in the outer AFC region (Fig. 5e) may be a sign of external stressors (Hetz et al., 2020; Wang et al., 2010). The *UPR* also orchestrates the enforcement of adaptive mechanisms to maintain an optimal rate of protein production and rapidly reacts to diverse stimuli, including extracellular responses to hormones, growth factors and small ligands that bind cell-surface receptors (Hetz *et al*., 2020). *UPR* is also a major contributor of the *CALR* marker for oAFC sub-populations (Fig. S2). *CALR* was previously found to be expressed in the oAF as well (Tam *et al*., 2020b) and it is a key gene in *UPR* pathway (Fig. S2). In addition to UPR signaling, the platelet-derived growth factor (*PDGF)* signaling also follows the increasing trend from inner to outer AF regions (Fig. 5f). *PDGF* expression has been shown to promote the proliferation of AFCs and NPCs (Pratsinis and Kletsas, 2007), suggesting a potential increased cell proliferation in the outer region. In short, the UPR and PDGF signaling may indicate an increased cell proliferation in response to external stimuli and stress applied to the outer AFCs. This theory was additionally supported by our detection of the expressed heat shock protein, *HSPA6,* in the outmost oAFC3 sub-population. Heat shock proteins (*HSP*s, Fig. 5a) were previously found to be strongly expressed in the outmost area of IVD and endplate cartilage in response to mechanical stress (Takao and Iwaki, 2002; Tam *et al*., 2020a). Collectively, the gradient trend of AFCs in neonatal IVD might be associated with certain biological function, for instance the elevated stress-induced cell proliferation and remodeling process among oAFCs. However, this needs to be confirmed by spatial single-cell analysis in the future.

The present work is not without limitations: our scRNA-seq lacks spatial resolution. The recently emerging single-cell spatial sequencing technology may be required in the next study.

In conclusion, we identified *FABP4*, *AKR1C1* and *APOE* as novel markers for human NC populations. We demonstrated the first direct comparison of human neonatal and adult IVD at a single-cell level, finding two distinct NC sub-populations in both neonatal and adult IVD. A pseudo- time trajectory analysis predicted the developmental process from neonatal IVD, composed of various cell types, into adult IVD, composed of mainly NPCs and AFCs. We also found that NCs preserve their cell identity, and some of their markers and function into adulthood. Lastly, we demonstrated a high heterogeneity of AFC populations in neonatal IVD and showed their gradual transitions of cell types, gene expression, function, signaling pathways, as well as their potential relevance in IVD development, which appears to be linked to a spatial gradient. Better understanding of these processes may help to identify the underlying factors contributing to age-associated diseases of the IVD and to develop tailored therapeutics that may delay or reverse these processes.

## STAR METHODS

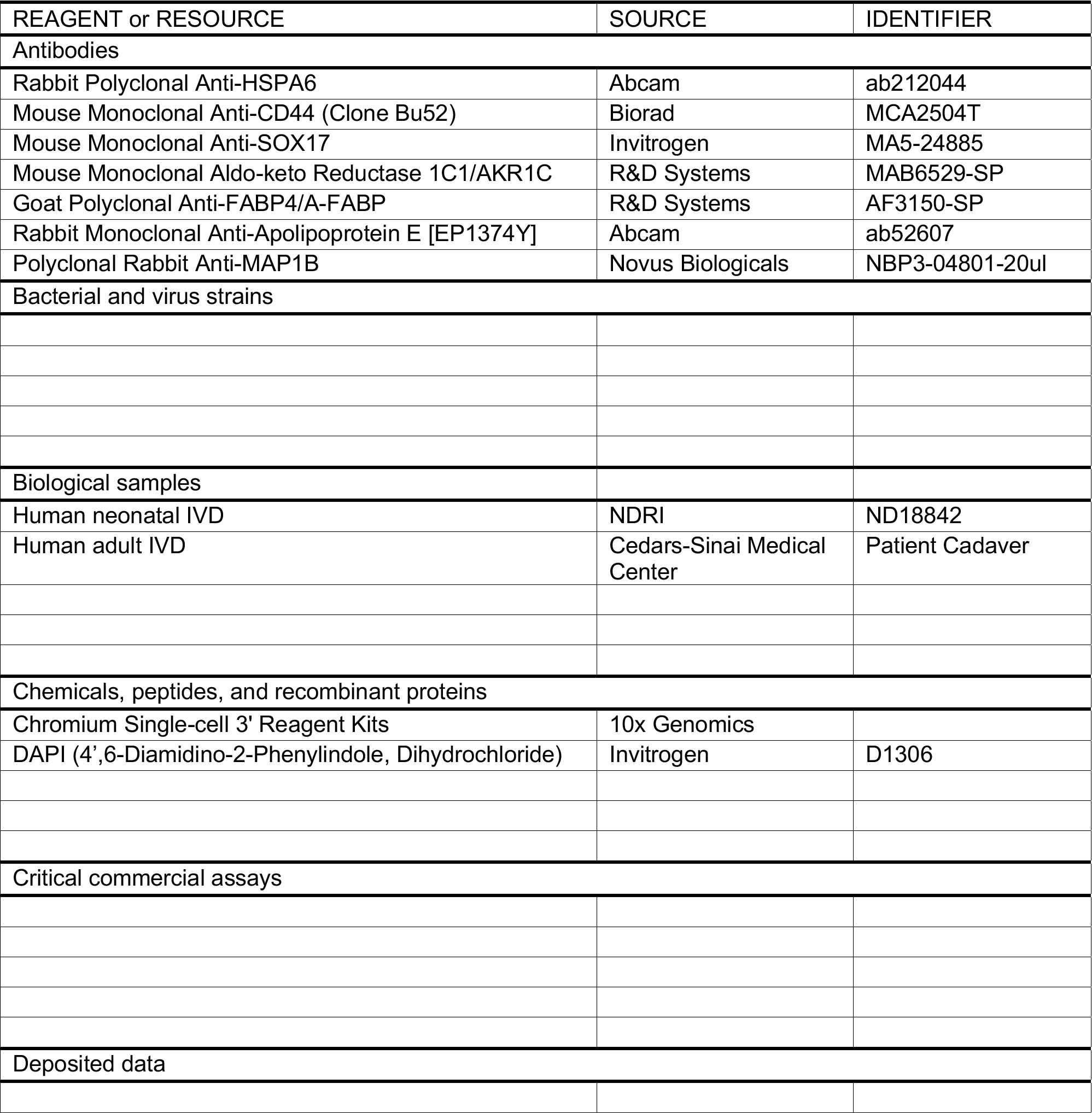
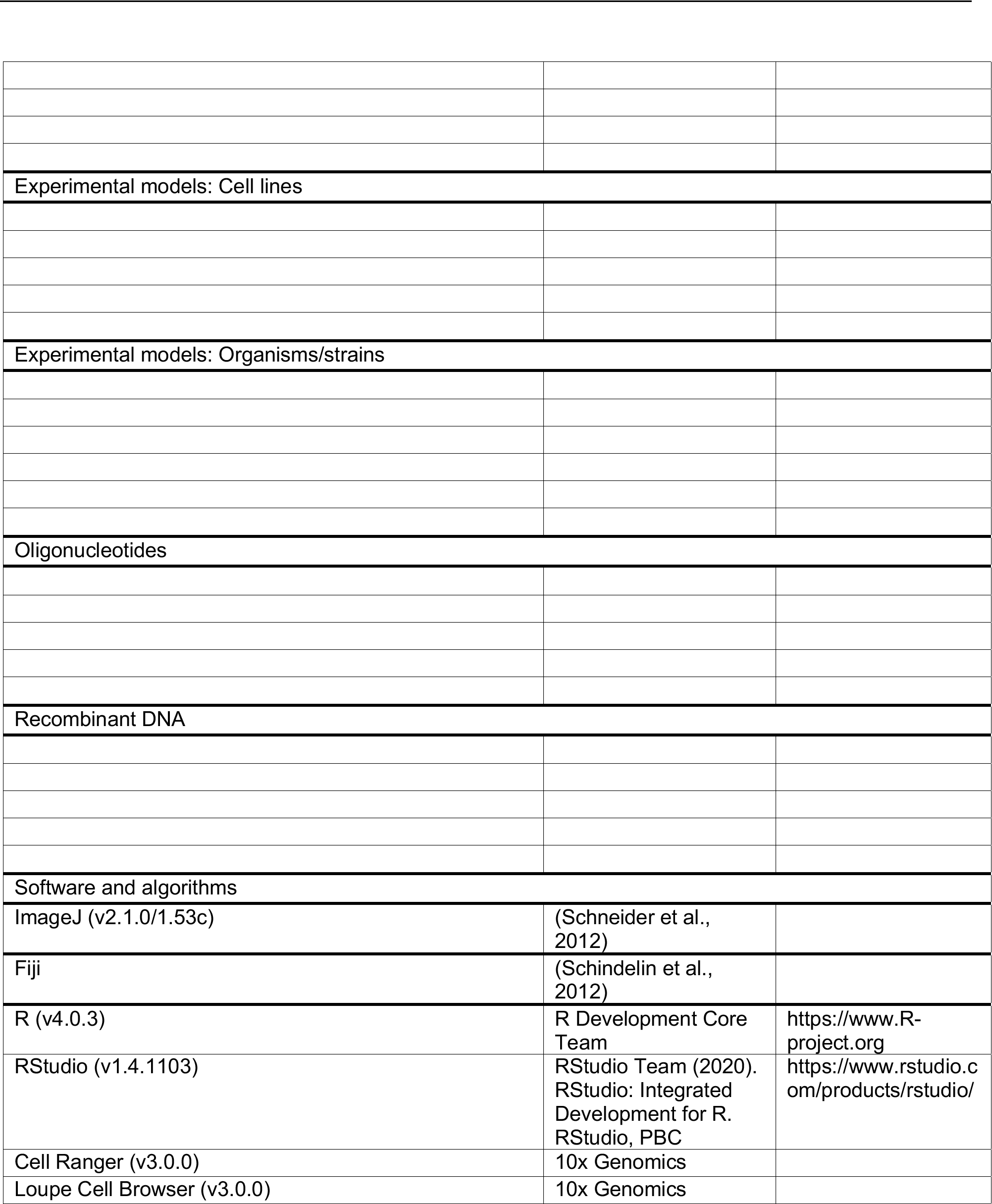
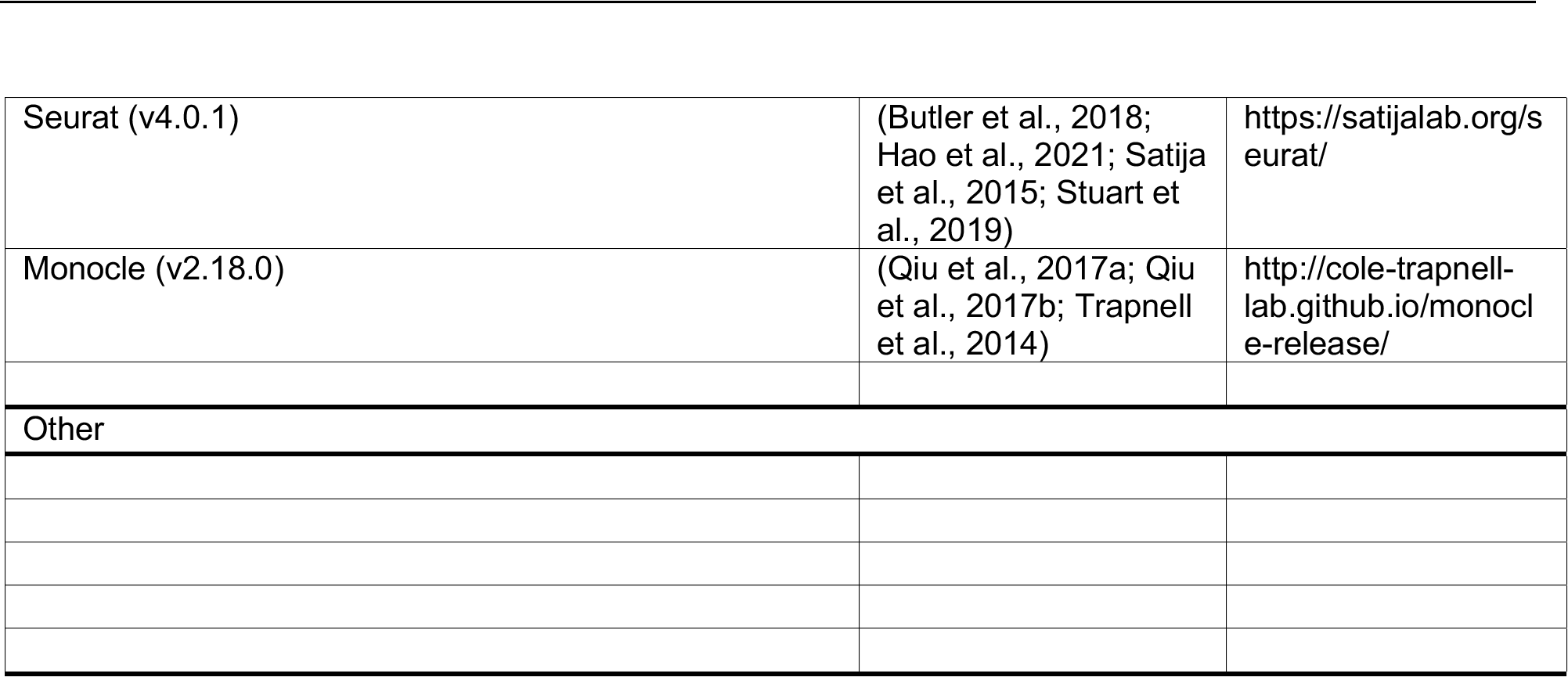
Key Resource Table

Key Resource Table
| REAGENT or RESOURCE | SOURCE | IDENTIFIER |
| --- | --- | --- |
| Antibodies |  |  |
| Rabbit Polyclonal Anti-HSPA6 | Abcam | ab212044 |
| Mouse Monoclonal Anti-CD44 (Clone Bu52) | Biorad | MCA2504T |
| Mouse Monoclonal Anti-SOX17 | Invitrogen | MA5-24885 |
| Mouse Monoclonal Aldo-keto Reductase 1C1/AKR1C | R&D Systems | MAB6529-SP |
| Goat Polyclonal Anti-FABP4/A-FABP | R&D Systems | AF3150-SP |
| Rabbit Monoclonal Anti-Apolipoprotein E [EP1374Y] | Abcam | ab52607 |
| Polyclonal Rabbit Anti-MAP1B | Novus Biologicals | NBP3-04801-20ul |
| Bacterial and virus strains |  |  |
| Biological samples |  |  |
| Human neonatal IVD | NDRI | ND18842 |
| Human adult IVD | Cedars-Sinai Medical Center | Patient Cadaver |
| Chemicals, peptides, and recombinant proteins |  |  |
| Chromium Single-cell 3' Reagent Kits | 10x Genomics |  |
| DAPI (4',6-Diamidino-2-Phenylindole, Dihydrochloride) | Invitrogen | D1306 |
| Critical commercial assays |  |  |
| Deposited data |  |  |
| Experimental models: Cell lines |  |  |
| Experimental models: Organisms/strains |  |  |
| Oligonucleotides |  |  |
| Recombinant DNA |  |  |
| Software and algorithms |  |  |
| ImageJ (v2.1.0/1.53c) | (Schneider et al., 2012) |  |
| Fiji | (Schindelin et al., 2012) |  |
| R (v4.0.3) | R Development Core Team | https://www.R-project.org |
| RStudio (v1.4.1103) | RStudio Team (2020). RStudio: Integrated Development for R. RStudio, PBC | https://www.rstudio.com/products/rstudio/ |
| Cell Ranger (v3.0.0) | 10x Genomics |  |
| Loupe Cell Browser (v3.0.0) | 10x Genomics |  |
| Seurat (v4.0.1) | (Butler et al., 2018; Hao et al., 2021; Satija et al., 2015; Stuart et al., 2019) | https://satijalab.org/seurat/ |
| Monocle (v2.18.0) | (Qiu et al., 2017a; Qiu et al., 2017b; Trapnell et al., 2014) | http://cole-trapnell-lab.github.io/monocle-release/ |
| Other |  |  |

## RESOURCE AVAILABILITY

### Lead contact

Further information and requests for resources and reagents should be directed to and will be fulfilled by the lead contact, Dmitriy Sheyn

### Materials availability

This study did not generate new unique reagents.

### Data and code availability

The raw sequencing data and original data are accessible in repositories.

## EXPERIMENTAL MODEL AND SUBJECT DETAILS

The Cedars-Sinai Medical Center Institutional Review Board approved this study (IRB number Pro00052234). Human neonatal and adult IVDs have both been investigated with scRNA-seq to reveal their cell atlas of respective age and to determine the difference between them. The neonatal samples were collected from 4 different spine levels (L1-L5) from a 6-hour postnatal male provided by the National Disease Research Interchange (NDRI, ND18842), 3 levels were used for the scRNAseq analysis and one for histology. The adult samples were collected from lumbar discs of 2 cadaveric spines and 1 clinical sample of removed and otherwise discarded disc. IVD tissues were harvested from three neonatal spinal disc levels and adult cadavers or clinical samples of patients with no back pain history (over 65 years old) were collected and analyzed via scRNA-seq (n=3 for neonatal IVDs, n=3 for adult IVDs). Additional lumbar levels were processed for histological analysis.

## METHOD DETAILS

### Single-cell Isolation and Sequencing

The IVDs were dissected and washed with Phosphate Buffered Saline (PBS). The tissue was manually minced to ∼1mm pieces in a sterile environment and enzymatically digested. Tissue was digested for 1 hour at 37°C in 2 mg/mL of Pronase Protease (Millipore, Temecula, CA) in growth media containing 1mM L-glutamine, 1% antibiotic-antimycotic solution (Gibco, Carlsbad, CA) and 10% fetal bovine serum (FBS, Gemini Bioproducts, West Sacramento, CA) in Dulbecco’s modified eagle media-F12 media (DMEM/F12, GIBCO, Carlsbad, CA). This was followed by ∼18 hours digestion at 37°C in 0.25 mg/mL of Collagenase Type 1S (Sigma Aldrich) in growth media. The resulting sample was pushed through a 70μm cell strainer and cells were isolated via centrifugation at 1200 rpm for ten minutes, The pellet was resuspended in phosphate buffered saline at a concentration of 1,500 cells/μL and 100uL of sample was processed for single-cell RNA sequencing using the Chromium Single-cell 3’ Reagent Kits from 10X Genomics.

Chromium Single-cell 3’ v2 libraries with ∼3,000 cells were prepared on a Chromium Controller with chips and reagents from Single-cell Gene Expression v2 kits following the manufacturer’s protocols (10x Genomics). The libraries were then sequenced using paired-end sequencing (28bp Read 1 libraries, and 91bp Read 2) with a single sample index (8bp) on an Illumina NovaSeq. Samples were sequenced to a depth of > 50,000 raw reads per cell, with raw sequencing data analyzed and visualized with pre-release versions of Cell Ranger 3.0.0 and Loupe Cell Browser 3.0.0. Raw sequencing data was demultiplexed and converted to fastq format by using bcl2fastq v2.20 (Illumina, San Diego, CA). Fragment analysis of indexed libraries was performed on the Agilent 4200 TapeStation (Agilent Technologies, Santa Clara, CA).

### Integration, Pre-processing, and Dimensional Reduction of the Single-cell Dataset of Neonatal and Adult IVD

Seurat package (v4.0.1) in R (v4.0.3) has been used to process the data of four samples. We firstly used *CreateSeuratObject* function to transform the loaded data into a Seurat object for each sample. Next, the four Seurat objects were combined into one matrix. *NormalizeData* and *FindVariableFeatures* were used. Afterwards, *FindIntegrationAnchors* and *IntegrateData* were used to integrate and anchor the 6 samples’ data together. *RunPCA* function was used to for principal component analysis with PC=30. *DimPlot* function was then used with “UMAP” reduction to obtain the 2D Uniform Manifold Approximation and Projection (UMAP) plot. *FindNeighbors* and *FindClusters* functions further classified all cells in 6 samples into 15 clusters at resolution of 0.5 (unsupervised clustering). The UMAP was then split by their sample type (neonatal or adult IVD) for comparison.

### Detection of Enriched Genes and Novel Markers for Notochordal Cells

The expression levels of NC relevant genes in neonatal or adult IVD were separately visualized using violin plot in log scale with split view. In the volcano plots, we set the cutoff threshold of FC to be > 2 and the cutoff threshold of adjusted *p* value to be < 1⨉10^-24^. Any genes falling out of this range were not considered as enriched genes or markers. Genes falling into this range were labeled with red color in the volcano plot. Next, top enriched genes from the red colored area were manually selected as novel *in silico* markers for specific cluster from neonatal IVD or adult IVD. Their expression levels were presented quantitatively in violin plots.

### Visualization of Single-Cell Atlases with 3D UMAP Plots

The cell coordination of the top three dimensions (PC1, PC2, and PC3) were extracted from the Seurat object integrating neonatal IVD and adult IVD. *FetchData* function was used to create the data frame for the 3D UMAP plot. The *plot_ly* function in *Plotly* package was used to obtain the final 3D UMAP plot. The color for each type was plot as the same as in 2D UMAP.

### Gene Ontology Term Enrichment and Pathway Analysis

Gene ontology (GO) term enrichment and pathway analysis was performed using Qiagen Ingenuity Pathway Analysis (IPA, version 65367011). A list of enriched genes for a specific cell sub-population (as compared against all other cells) was firstly generated in R studio using *FindMarkers* function. The specific cell sub-populations include NC1-2, iAFC, oAFC1-3 respectively. In other attempts, we did multiple further analyses: we created enriched gene list for NC by combining NC1 and NC2 cell sub-populations followed by *FindMarkers* function against all other cells; we created enriched gene list for NPC by combining NPC1-4 cell sub-populations followed by *FindMarkers* function against all other cells. All the above enriched gene lists were generated for both neonatal IVD and adult IVD. The adjusted *p* value and fold change for genes with *p*<0.0001 were loaded into the IPA software. We included the database for human tissue and primary cells but excluded those for cell lines, as our samples were harvested from human samples only. The analyses covered canonical pathways, upstream analysis, diseases and biological functions, and regulatory networks. The heatmaps for enriched biological functions, enriched pathways, and key genes in the pathways were replotted using GraphPad Prism (v9.2.0). We demonstrated the most relevant results in figures.

The comparison of gene overlap in NCs and NPCs in both neonatal and adult IVD was performed at Metascape.org using Circos plot. The enriched genes with *p*<0.0001 and FC>1 were selected for Circos plot analysis.

### Histology and Immunofluorescence

IVD samples from the sample were fixed in 10% phosphate buffered formalin for three days, decalcified by incubation in 0.5M EDTA (pH 7.4, Sigma Aldrich) for three weeks, and embedded in paraffin. Five-micron-thick sections were cut from the paraffin blocks. Hematoxylin and eosin (H&E) staining was performed to evaluate the morphological features. Images were captured using a Carl Zeiss Axio Imager Z1 fluorescent microscope (Carl Zeiss, Germany) equipped with ApoTome and AxioCam HRc cameras. Images were analyzed using QuPath software (v0.2.3).

For immunofluorescence, sections were deparaffinized, rehydrated in DPBS, treated with antigen retrieval solution at room temperature for 20 minutes (Dako # S3020, Agilent Technologies). Slides were washed with PBS, blocked with 3% normal donkey serum (Jackson ImmunoResearch Laboratories Inc., West Grove, PA) in 0.3% Triton-X (Sigma Aldrich, St. Louis, MO) and again washed with PBS. Slides were stained with primary antibodies (anti-human *HSPA6, CD44, SOX17, AKR1C1, FABP4, APOE*). The primary antibodies were applied to the slides, after which the slides were incubated at 4°C overnight and washed using PBS; the slides were then incubated with secondary antibodies for 1hr at room temperature. Finally, the slides were stained with 4’,6- diamidino-2-phenylindole dihydrochloride (DAPI, 0.28 µg/ml) for 5 min in the dark. Subsequently, sections were washed three times in dPBS and mounted with Prolong Gold with DAPI (Life Technologies).

### Pseudo-Time Analysis of Developmental Trajectories

The pseudo-time developmental trajectory analysis was performed using *Monocle* package (v2.18.0). The original integrated Seurat object was subset into a new Seurat object that contains the following cell populations: NC (as the combination of NC1-2), NPC (as the combination of NPC1-4), iAFC, and oAFC (as the combination of oAFC1-3). Low-quality cells were filtered by the following standards: 1) Cells whose total mRNAs >10^6^ were filtered; 2) Cells whose total mRNAs were out of a range of mean ± 2 ! standard deviations were filtered. The gene marker was also filtered out when less than 10 cells expressed this maker. The data matrix was then log-transformed and standardized to the same scale. We chose up to 500 top globally enriched genes for each sub-populations to determine cell progress. The dimensions of the dataset were reduced using DDRTree method. The trajectories were plotted using *plot_cell_trajectory* function. The cells in trajectories were colored with either their ages, cell types, or pseudo-time states.

## ACKNOWLEDGMENTS

This study was supported by NIH/NIAMS, Grant/Award Number: K01AR071512. The authors wish to acknowledge the Cedars-Sinai Genomics Core for next generation services and the Cedars-Sinai Biobank for human adult cadaver sample. The authors thank National Disease Research Interchange (NDRI) for human neonatal sample and Mrs. Julia Sheyn for the fluorescent images acquisition.

## AUTHORS CONTRIBUTIONS

Conceptualization, W.J., D.S., J.G., and R.H.; Methodology, W.J., R.H., D.S., P.M., and A.W.; Software, W.J., R.H., and A.W.; Validation, K.S and G.K.; Formal Analysis, W.J., R.H., and D.S.; Investigation, W.J., D.S., and R.H.; Resources, R.H., D.S. and W.J.; Data Curation, W.J. and R.H.; Writing – Original Draft, W.J.; Writing Review & Editing, J.G., D.S., W.J., R.H., and G.K.; Visualization, W.J., D.S., and R.H.; Supervision, D.S., and R.H.; Funding Acquisition, D.S.

---

Figure S1. Additional Pathway and Network Analyses for NC Sub-Populations and Key Regulators in NC Populations Compared with NPC Populations in Neonatal and Adult IVD. Pathway and network summary for a) NC1 and b) NC2 sub-populations in neonatal IVD. c) The IL1-regulated network involving *APOE* and *AKR1C1/AKR1C2.* d) Heatmap showing top regulators for NCs and NPCs in neonatal and adult IVD. Orange indicates activation, blue deactivation, and grey shows data that were either not detected or did not pass the filtering.

Figure S2. Gene Expression Projected on the Spatially Dependent Unfolded Protein Response Signaling Pathway for oAFC2 Sub-Populations in Neonatal IVD.

Figure S3. Comparison of the Spatially Dependent Trend of Expression Levels for Collagen-Relevant Genes in Neonatal and Adult IVD.

Figure S4. Heatmap Showing the Transcriptomes of Top Differentially Expressed Genes for All Sub-Populations Identified in Human IVD. Data include both neonatal and adult IVD.

