## Supplemental Figures for "Single-cell Atlas Unveils Cellular Heterogeneity and Novel Markers in Human Neonatal and Adult Intervertebral Discs"

### Supplemental Information. Figure S1

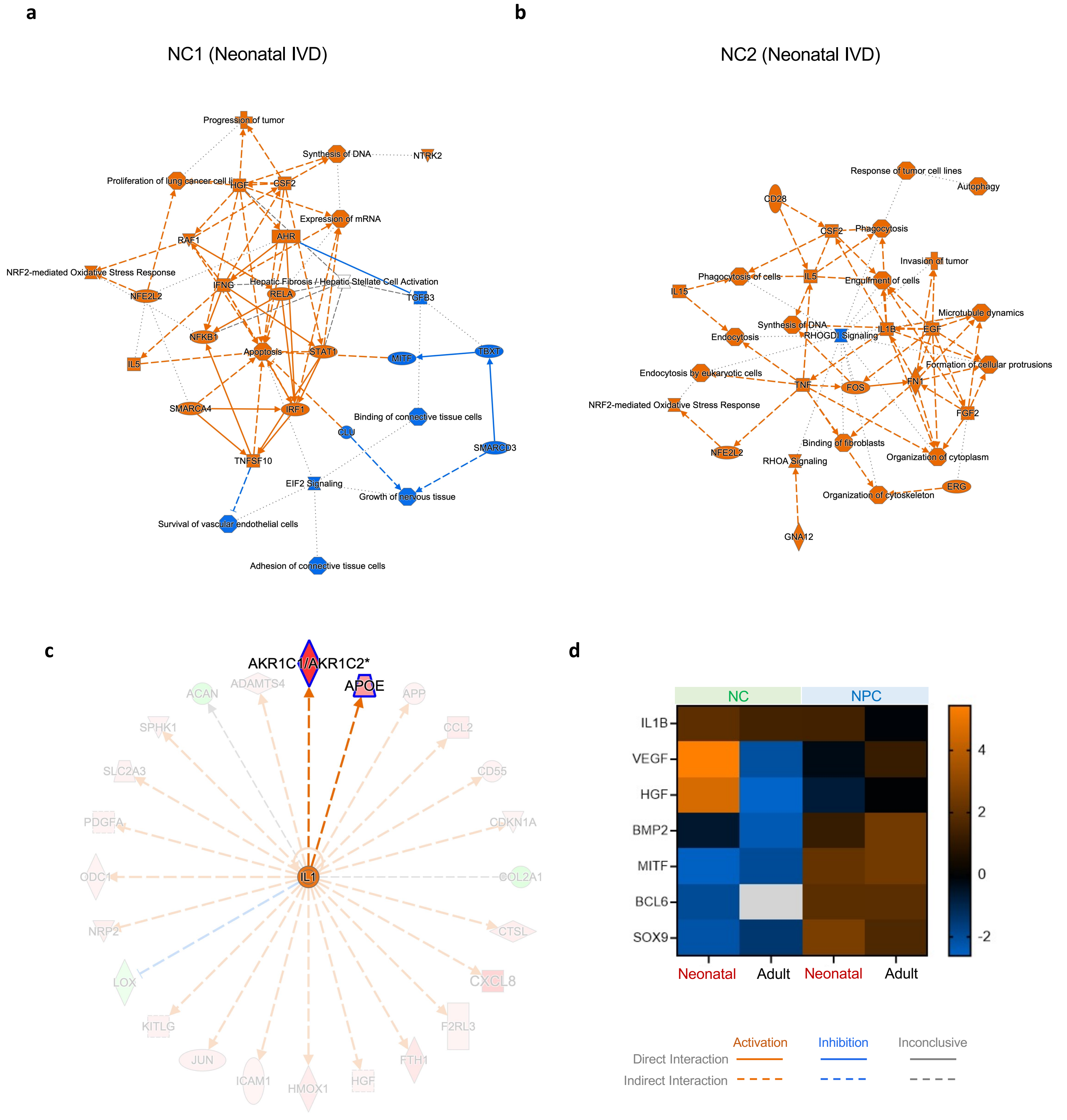


**Figure S1. Additional Pathway and Network Analyses for NC Sub-Populations and Key Regulators in NC Populations Compared with NPC Populations in Neonatal and Adult IVD.** Pathway and network summary for **a)** NC1 and **b)** NC2 sub-populations in neonatal IVD. **c)** The IL1-regulated network involving *APOE* and *AKR1C1/AKR1C2.* **d)** Heatmap showing top regulators for NCs and NPCs in neonatal and adult IVD. Orange indicates activation, blue deactivation, and grey shows data that were either not detected or did not pass the filtering.

### Supplemental Information. Figure S2

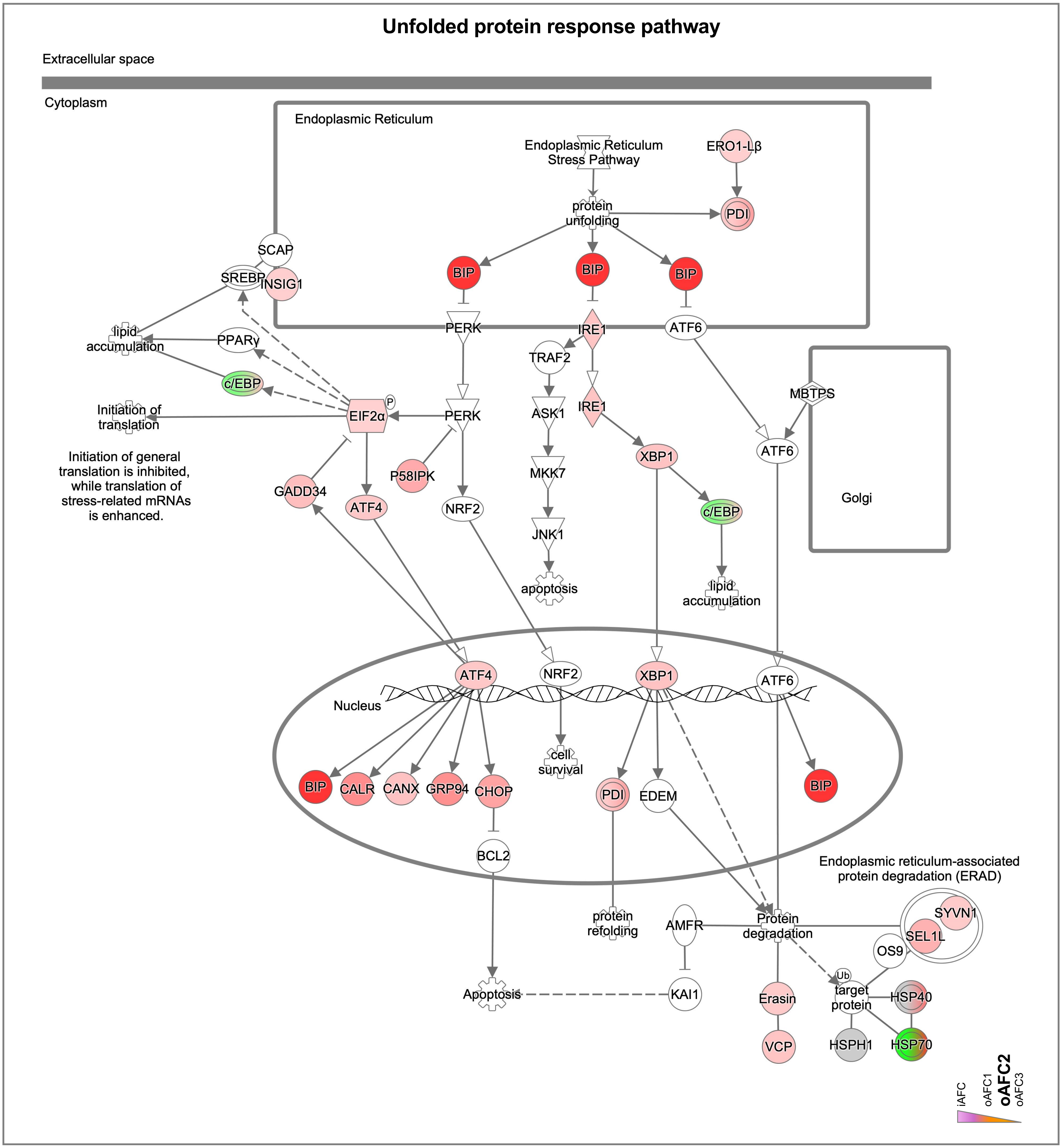


**Figure S2. Gene Expression Projected on the Spatially Dependent Unfolded Protein Response Signaling Pathway for oAFC2 Sub-Populations in Neonatal IVD.**

### Supplemental Information. Figure S3

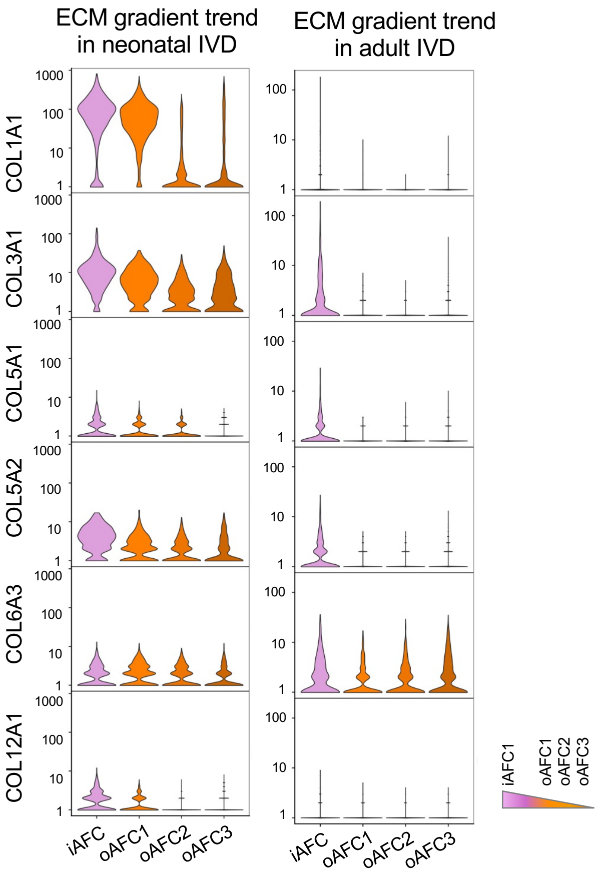


**Figure S3. Comparison of the Spatially Dependent Trend of Expression Levels for Collagen-Relevant Genes in Neonatal and Adult IVD.**

Supplemental Information. Figure S4

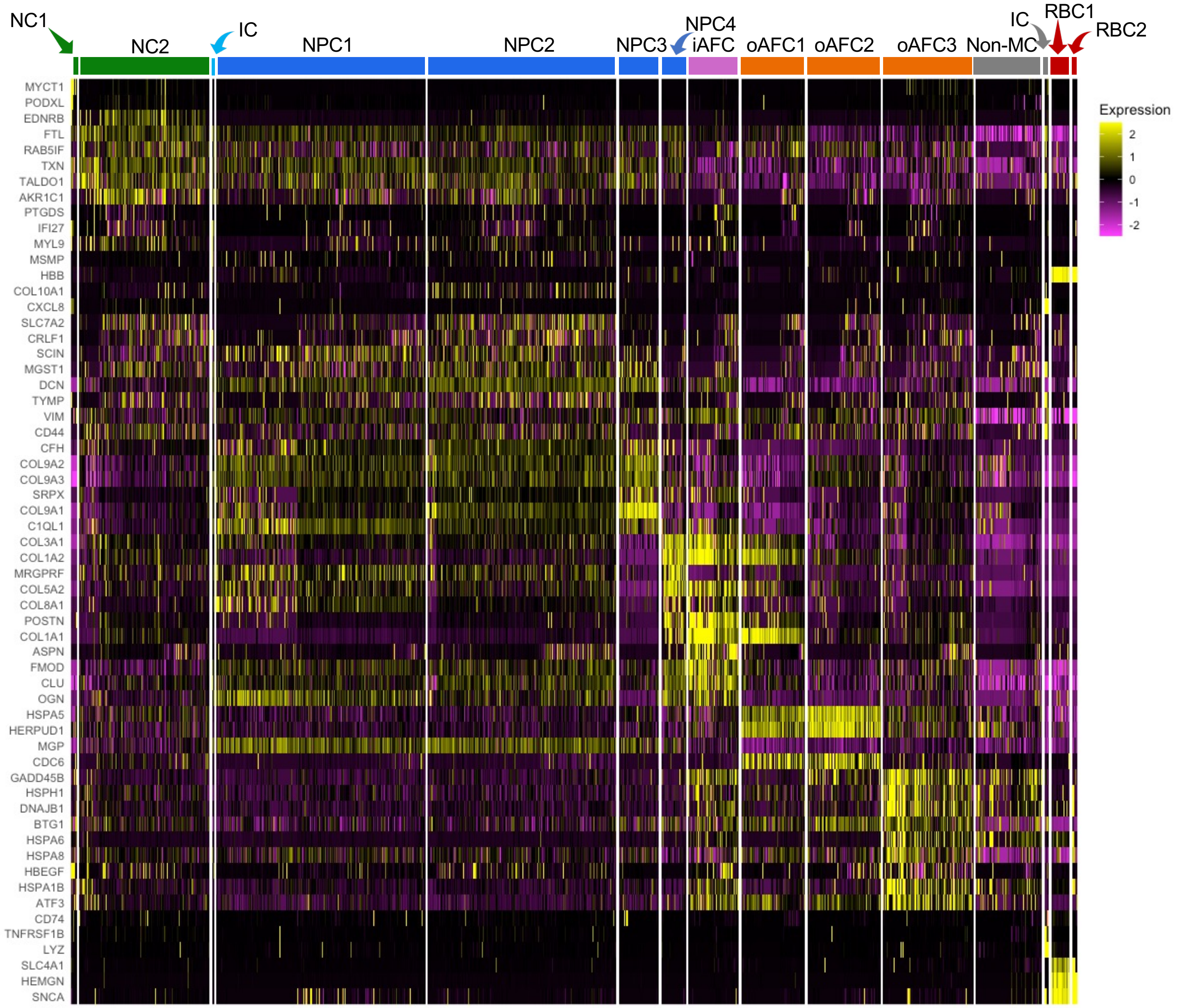


**Figure S4. Heatmap Showing the Transcriptomes of Top Differentially Expressed Genes for All Sub-Populations Identified in Human IVD.** Data include both neonatal and adult IVD.
